# Importance of environmental signals for proper cardiac morphological development in Atlantic salmon

**DOI:** 10.1101/2024.02.19.580955

**Authors:** Marco A. Vindas, Vilde Arntzen Engdal, Simona Kavaliauskiene, Ole Folkedal, Erik Höglund, Marta Moyano, Øyvind Øverli, Michael Frisk, Ida B. Johansen

## Abstract

The hearts of salmonids display remarkable plasticity, adapting to various environmental factors that influence cardiac function and demand. For instance, in response to cold temperature, the salmonid heart undergoes growth and remodeling to counterbalance the reduced contractile function associated with dropping temperatures. Alongside heart size, the distinct pyramidal shape of the wild salmonid heart is essential for optimal cardiac performance, yet the environmental drivers behind this optimal cardiac morphology remain to be fully understood. Intriguingly, farmed salmonids often have rounded, asymmetrical ventricles and misaligned bulbi from an early age. These deformities are noteworthy given that farmed salmon are often not exposed to natural cues, such as a gradual temperature increase and changing day lengths, during critical developmental stages. In this study, we investigated whether natural environmental conditions during early life stages are pivotal for proper cardiac morphology. Atlantic salmon were raised under simulated natural conditions (low temperature with a natural photoperiod; SIMNAT) and compared with those reared under simulated farming conditions (SIMFARM). Our findings reveal that the ventricle shape and bulbus alignment in SIMNAT fish closely resemble those of wild salmon, while functional analyses indicate significant differences between SIMNAT and SIMFARM hearts, suggesting diastolic dysfunction and higher cardiac workload in SIMFARM hearts. These findings highlight the profound influence of environmental factors such as water temperature and photoperiod on the structural development of the salmonid heart, underscoring the importance of early environmental conditions for cardiac health.

## Introduction

Animals that undergo dramatic habitat shifts during their life cycle are characterized by complex adaptations which helps in their survival. This includes physiological adaptations of vital organ systems that are synchronized by environmental signals, such as day length and temperature (McCormick, 2012; West and Wood, 2018). The Atlantic salmon *(Salmo salar)* has a complex life cycle, with spawning and juvenile stages taking place in fresh water, before migration to sea water to grow and mature (Stefansson et al., 2008). Furthermore, salmon live in cold temperate areas where they remain active at a range of temperatures (Jonsson and Jonsson, 2009). Thus, salmon needs to adjust to both variation in salinity and temperature and has evolved complex physiological adaptations allowing such a lifestyle (McCormick, 2012). For example, the salmonid heart grows and remodels in response to cold temperatures to compensate for the negative effect on decreasing temperature on contractile function (Graham and Farrell, 1989; Keen et al., 2017). Similarly, the transition from freshwater to sea water is associated with myocardial growth (compactum) which generates increased stroke volumes and cardiac output (Brijs et al., 2017). This cardiovascular response is believed to compensate for hemodynamic changes associated with increased salinity, as well as an increased demand for circulation of osmoregulating organs (Brijs et al., 2015). In this context, there are indications that cardiac preparations for sea water transition are initiated already when salmonids go through their metamorphosis for seawater adaptation (*i.e.* smoltification). In wild salmonids, smoltification is triggered by photoperiod and temperature cues (*i.e.* gradual increase in day length and water temperature) and involve a range of behavioral, physiological, and morphological changes preparing the fish for downstream migration and sea water entry (Stefansson et al., 2008). For example, relative heart size and the compact to spongy muscle ratio increase from the parr to the smolt stage, when fish transition to a more pelagic lifestyle (Graham and Farrell, 1992; McCormick, 2012). However, the specific environmental triggers and physiological mechanisms underlying this cardiac remodeling remain largely unknown.

While heart size plays a crucial role, heart shape is equally pivotal for optimizing cardiac and physical performance (Claireaux et al., 2005). This is evident in certain athletic fish species, such as salmonids and tunas, that exhibit a distinct heart morphology characterized by an elongated pyramidally shaped ventricle (Anttila and Farrell, 2011). Unlike hypertrophic remodeling, the factors driving morphological alterations and development of this geometrically advantageous heart shape remain poorly understood.

Interestingly, farmed salmon exhibit a substantially different heart shape than their wild counterparts. Farmed salmon display rounded and asymmetrical ventricles, misaligned bulbi, and a notable accumulation of fat deposits on both bulbus and ventricle (Brijs et al., 2020; Frisk et al., 2020; Handeland et al., 2003). Recent research revealed that some of these distinct morphological differences between wild and farmed salmon manifest in the early freshwater stage, while others become apparent closer to smoltification (Frisk et al., 2020). Thus,environmental factors during early life, from hatching to smoltification, emerge as potential key influencers of cardiac remodeling and development.

In this context, farmed salmon serve as valuable models for investigating the role of environmental factors on cardiac remodeling. They are often deprived of natural environmental cues and reared under conditions that diverge significantly from the natural environment experienced by their wild counterparts. Commercial Atlantic salmon hatcheries, for instance, typically rear salmon under continuous light and at elevated temperatures to promote accelerated growth during the freshwater stage (Handeland et al., 2003; Steffanson et al., 1991). Notably, two recent studies provide evidence that cardiac morphological deviations in farmed salmonids are associated with elevated temperatures during early rearing(Brijs et al., 2020; Frisk et al., 2020). For example, Brijs et al. (Brijs et al., 2020) showed that rainbow trout exposed to warm temperatures (10 to 15 °C) with light manipulation (19:5-h light:dark) in the hatchery displayed more severe ventricular morphological deviations compared to rainbow trout reared at more natural conditions. Similarly, Frisk et al. (Frisk et al., 2020) linked elevated temperature in the hatchery to more severe ventricular morphological deviations later in life in Atlantic salmon. Thus, common cardiac deviations frequently observed in aquaculture may represent maladaptive remodeling of the heart in response to unfavorable rearing conditions early in life, but this remains to be explored.

Here, we test the hypothesis that exposure to a more natural environmental regime during early rearing stages is necessary for proper morphological development of the heart in Atlantic salmon. We reared Atlantic salmon under either simulated natural conditions (*i.e.* low temperature and a simulated natural photoperiod; SIMNAT) or simulated farmed conditions (*i.e.* elevated temperature and periods of continuous photoperiod; SIMFARM) conditions and compared their heart morphology throughout the experimental period. Heart morphology was compared to wild Atlantic salmon fry, parr and smolt. Overall, SIMNAT fish exhibited a closer resemblance to wild salmon than SIMFARM hearts, clearly indicating that environmental signals, like water temperature and photoperiod, play significant roles in the geometric modelling of the salmonid heart. Importantly, functional analyses uncovered significant differences between SIMFARM and SIMNAT hearts, suggesting the adverse effects of altered morphology on cardiac health in SIMFARM fish.

## Materials and methods

### Ethics Statement

The experiments were performed in accordance with current Norwegian law for experimentation and procedures on live animals and were approved by the Norwegian Food Safety Authority under permit number 24848.

### Fish husbandry

The fish used in this experiment were from the AquaGen Atlantic QTL-innOva SHIELD strain. A total of 3600 fish were hatched in two batches (*n* = 1800 per treatment) on site at the Matre Research Station facilities, Institute of Marine Research, Norway. The aim of the experiment was to compare how rearing conditions affect heart morphology by coordinating a common smoltification for fish reared under simulated natural (SIMNAT) and simulated farmed (SIMFARM) conditions. Since simulated natural conditions are associated with slower growth than the simulated farmed conditions, the SIMNAT group batch was hatched in December 2019, while the SIMFARM group batch was hatched ten months later (September 2020) which resulted in both groups smoltifying in May 2021. During egg incubation, the conditions were the same for both groups (8 °C and total darkness), but from hatching to start-feeding the SIMNAT group was kept at 10 °C and the SIMFARM at 13 °C, both at 24 h light. After start-feeding, the SIMNAT group was kept at 7 °C and on a natural photoperiod until smoltification in May 2021. Meanwhile, the SIMFARM group was kept at 13 °C and at 24 h light for six months, followed by a 12:12 light/dark period (winter signal) before returning to a 24-light regime until smoltification in May 2021. At the onset of the winter signal, temperature was also decreased to 10 °C for the SIMFARM fish to avoid extreme size disparities between the groups. Both treatment groups were kept on a continuous flow of filtered, UV-C treated, and aerated water, which ensured oxygen saturation >85% levels at all times and prevented waste products from accumulating. Triplicates for each treatment were maintained until smoltification. Circular 100 L tanks were used until the fish reached 1.54 g at which point, they were transferred to square 500 L tanks until smoltification. The fish were fed to satiation daily with commercial pellets (Nutra Olympic, following growth tables by Skretting), between 14:00 and 18:00 via automatic feeding devices. All fish were vaccinated after reaching approximately 30 g with AQUAVAC® PD7 (MSD Animal Health) following standard salmon aquaculture procedures. See Figure 1 for a schematic illustration of the experimental design.

**Figure 1.**
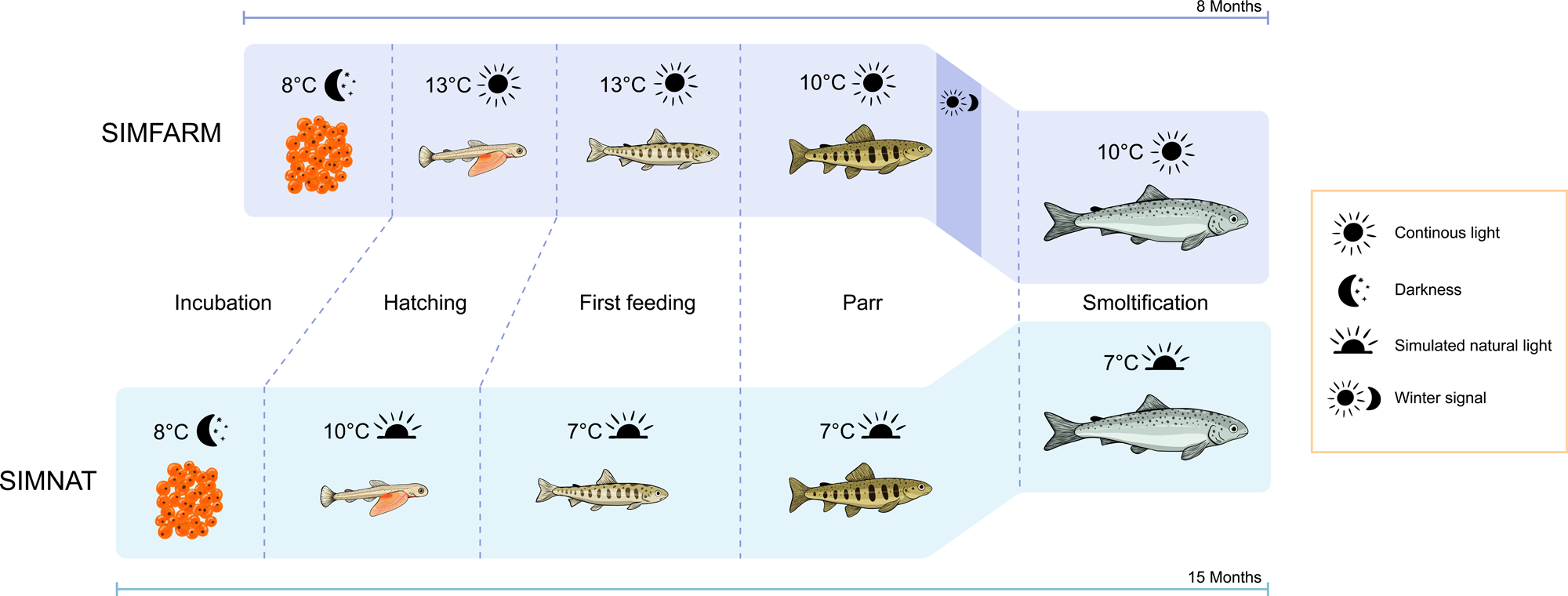
Schematic representation of the experimental design timeline for the Atlantic salmon groups reared in simulated natural (SIMNAT) and simulated farmed (SIMFARM) conditions.

### Sampling

The treatment groups were sampled at four time points throughout the experimental period: (1) at the fry stage, (2) during the parr stage (both experimental groups were sampled at the onset of the winter signal for the SIMFARM group), (3) at the estimated start of smoltification, and (4) at the end of the smoltification (fish for this sampling were sampled at 3 different occasions during one week, but the data was pooled since they all represent the same timepoint). At all sampling occasions, four fish were randomly chosen from each tank for a total of 12 fish per timepoint/treatment, except for the first timepoint (the fry stage point) at which 15 fish were sampled (five per tank). At all samplings, fish were netted from their home tanks and immediately euthanized with a lethal dose of buffered MS-222 at a concentration of 2 g/L until completely unresponsive and motionless (within approximately 30 s). Fish were rapidly weighed, fork length was measured, and a blood sample was taken from the caudal vein using a 23G, 1 ml syringe containing the anticoagulant ethylene diamine tetra acetic acid (EDTA). Following centrifugation for 10 min at 9289 rcf and 4 °C, plasma samples were frozen and stored at −80 °C for later analysis. The hearts were then removed and stored at 4 °C in buffered 10% formalin solution.

### Wild fish sampling

Atlantic salmon fry and parr were captured using electrofishing gear in Hagnesvassdraget, a western tributary of Numedalslågen in Kvelde, Norway (59.19753° N, 10.10184° E) in August 2021. A total of 24 fry (1–7 g) and 28 parr (5–45 g) were euthanized and sampled as explained above.

Downstream migrating wild Atlantic salmon smolts were caught at two points in the River Nidelva, Agder, Norway (58.41540° N, 8.74242° E) in May 2021. First, smolts were collected using a modified fyke net placed in the Songeelva river, a tributary of the Nidelva river, 25 km upstream of the river mouth. Second, fish were caught in a Wolf trap located at the entrance of the surface fish passage at Rygene hydropower plant, 9.4 km downstream of the river mouth. In both cases, fish were collected from the trap and transported to collection tanks at the Rygene hydropower plant. Fish were euthanized and sampled as explained above.

### Imaging and analysis of cardiac morphology

Fixated hearts were photographed from the left lateral and ventrodorsal projections inside a Styrofoam box (Internal H:24cm W: 21 cm L: 27 cm) lit by an internal LED-light (Northlight LEDlamp, Art. No. 36-6465). Photographs were captured with a Cannon EOS 4000D camera with an EF-S 18-55 III lens mounted on the Styrofoam box. Thus, all hearts were photographed from the same angles and under standardized light conditions. All heart measurements were calculated and analyzed according to Engdal et al. (Engdal et al., 2024). Specifically, the ventricular height:width ratio was analyzed from the ventrodorsal projection by dividing the ventricle length (from apex to bulbus) by the ventricle width (widest part of ventricle). In the left lateral projection, ventricular symmetry was quantified by measuring the angle between the ventricular vertical axis and the axis running from the ventriculobulbar groove to the left dorsal ventricular apex. In the same projection, alignment of bulbus arteriosus was quantified by measuring the angle between the bulbular horizontal axis and the ventricular vertical axis. Finally, the bulbus width:ventricular width was quantified from the ventrodorsal projection by dividing the bulbus width (widest part of the bulbus) by the ventricle width (widest part of the ventricle). All photos were analyzed using the Fiji platform (Schindelin et al., 2012) by ImageJ2 (Rueden et al., 2017).

### Magnetic resonance imaging (MRI) of fixed ventricles

Fixed hearts were placed in Dulbecco’s phosphate buffered saline (DPBS) with 0.5 mol/L Magnevist® (Bayer Schering Pharma, Berlin, Germany) for 1-2 days. Then, the hearts were dried with paper tissue and filled with Fomblin® Y LVAC 06/6 (Solvay Specialty Polymers Italy SpA, Bollate, Italy) through the atrium prior to mounting in 15 ml plastic tubes using cotton soaked in Fomblin®. MRI scans were performed on a 9.4T MRI system equipped with Avance Neo console and Paravision 360 software (Bruker Biospin, Germany) and a 19 mm quadrature-driven birdcage RF coil (Rapid Biomedical, Germany). A 3D gradient echo (FLASH) sequence was used for acquisition with a XYZ resolution of 25 x 25 x 200 μm, TR/TE of 35/6 ms, flip angle of 30 degrees, and averaging of 12.

### MRI image analysis

For compactum thickness analysis, five ventricular sections were selected: the section at the widest part of the ventricle, and four additional sections toward the apex with 5% increment of the total ventricular length. This resulted in 5 images within 20% of the total length of the ventricle (Supplementary Figure 1). Image analyses were performed using the Fiji platform. The total ventricular area was semi-automatically selected using the “magic wand” function. Any irregularities due to the presence of air bubbles or tearing of the tissue were manually removed prior to further analysis. The spongiosum was manually selected, and the compactum was then selected as the difference between the whole ventricle and the spongiosum. Compactum thickness was measured using the “Local Thickness” function in Fiji. The resulting thickness map was inspected manually for any irregularities, and mean thickness of compactum was measured as an average of all pixels in the thickness map. To adjust for differences in ventricle size between hearts, compactum thickness was normalized to the square root of the spongious area in each section. The normalization is based on the approximation of the spherical geometry of the heart, and the assumptions that pressure and wall stress do not depend on heart size (Norton, 2001).

Volume analysis of different heart compartments was performed using a 3D Slicer 5.2.1 software (Fedorov et al., 2012). Whole heart segmentation was performed using automatic thresholding (Otsu method) followed by manual inspection for air bubbles and selection of the lumen. Both the bulbus and atrium were manually removed from the selection to obtain ventricular measurements only. The ventricular lumen segment was defined as the difference between filled and empty (after automatic thresholding) ventricular segments. Segment statistics were automatically obtained using a built-in function in the 3D Slicer.

### Echocardiography

The functional effects of altered heart morphology were examined in a subset of SIMFARM (n = 10) and SIMNAT (n = 10) individuals by echocardiography (Supplementary Figure 2). These individuals are part of a subgroup of fish that were kept in their home tanks under the same described conditions, except for water salinity. That is, the tanks were switched to seawater (33 ppt) 48 h before echocardiography measurements. Fish were anesthetized one by one, in 150 mg/L MS-222 (Sigma-Aldrich). When locomotory and gill movements ceased, individuals were transferred to a V-board where gills were irrigated with temperature-controlled water (10 °C) containing 75 mg/L MS-222 of anesthesia (to maintain light sedation). Echocardiography was performed with a linear probe (GE 12L-RS at 13MHz) connected to a Vivid iq ultrasound system (GE Vingmed Ultrasound A/S, Horten, Norway). The probe was placed along the fishes’ longitudinal axis using the atrio-ventricular (AV) valve, ventricular dorsal apex, and ventro-bulbar (VB) valve of the heart as reference points for standardized imaging projections. Cardiac structure was visualized with b-mode, and hemodynamics (AV and VB blood flow velocities) were recorded in pulsed-wave Doppler mode. Every single recording lasted for at least three heartbeats and the whole imaging protocol took less than 10 minutes per fish. Images were analyzed with EchoPAC v. 203 (GE Vingmed), and all parameters presented are an average of three heartbeats from each individual. Information about systolic function was obtained by assessing outflow tract diameter (b-mode) and outflow tract blood flow velocities with pulsed wave (PW) Doppler. These measurements were used to calculate the outflow tract pressure gradients and the blood velocity time integral to allow estimation of cardiac output using a built-in function in EchoPAC. Early early and late diastolic atrio-ventricular valve velocities (E- and A-wave, respectively) as well as their ratio (E:A) were measured during ventricular filling to obtain information about diastolic function. Analyses of all echocardiography parameters, except cardiac output and stroke volume, were performed using built-in algorithms in EchoPAC (GE Healthcare). Due to the distinct hemodynamic pattern in the salmonid outflow tract cardiac output and stroke volume were calculated as described by Ma *et al*.(Ma et al., 2019).

### Statistical analysis

RStudio software 4.0.4 (R Development Core Team, http://www.rproject.org) was used for the statistical analyses. The statistical packages’ nlme’ and ‘MuMIn’ were used for exploratory linear mixed effect models (LME). The heart morphology measurements (including volumes and thickness) and biometric data were analyzed using an LME, with treatment (Wild *vs.* SIMNAT *vs.* SIMFARM) and time (fry stage, parr, and end of the smoltification period) as categorical independent variables or treatment (SIMNAT *vs.* SIMFARM) and time (fry stage, parr stage, start of the smoltification and end of the smoltification) as categorical independent variables with both LMEs including tank as a random effect for the experimental groups. In addition, the surface area and volume MRI measurements at smoltification were analyzed using an LME, with treatment (Wild *vs.* SIMNAT *vs.* SIMFARM) as a categorical independent variable and fish as a random effect. The initial LME models allowed the independent variables to interact. However, the final model was selected based on the lowest Akaike information criterion (AICc) score, *i.e*., the model with the best data fit when weighted against model complexity. Visual inspection of the qqnorm and residual plots to check the assumptions of normality and homoscedasticity confirmed that these models conformed to these assumptions. Interactive effects between treatment and test were assessed using Tukey–Kramer honestly significant differences post hoc test. Functional echochardiography data were analyzed with a Student’s t-test between SIMFARM and NATFARM. A correlation matrix using the R package HMSC with Benjamini-Hochberg *p* value correction, was used to explore interactions between CSI, morphological measurements, and functional echocardiography data. Significance was assessed as p ≤ 0.05.

## Results

### Biometric differences between SIMFARM, SIMNAT and wild salmon

Both treatment (W: χ*^2^_(1)_* = 11.7, *p* < 0.001 and L: χ*^2^_(1)_* = 16, *p* < 0.001) and time (W: χ*^2^_(1)_* = 265, *p* < 0.001 and L: χ*^2^_(1)_* = 2182, *p* < 0.001) influenced body mass (W) and body length (L) throughout the experiment. Whereas both groups grew heavier and longer with time, the SIMFARM fish were consistently heavier and longer than the SIMNAT groups throughout the experimental period (Table 1).

**Table 1.** Mean ± SEM body mass (W; in g) and length (L; in cm) throughout the experiment for Atlantic salmon reared under simulated natural (SIMNAT) and farmed conditions (SIMFARM).

| Treatment | Fry stage | Parr stage | Start smoltification | End smoltification |
| --- | --- | --- | --- | --- |
| SIMFARM | $2 \pm 0.1$ W | $43.7 \pm 2.5$ W | $56.3 \pm 2.4$ W | $74.4 \pm 2.2$ W |
| | $6.6 \pm 0.1$ L | $14.9 \pm 0.3$ L | $16.1 \pm 0.2$ L | $18.7 \pm 0.2$ L |
| | | | | $0.169 \pm 0.003$ CSI |
| | | | | $0.09 \pm 0.003$ RVM |
| SIMNAT | $1.3 \pm 0.1$ W | $37.1 \pm 2.3$ W | $50.7 \pm 3$ W | $65.7 \pm 2.5$ W |
| | $4.9 \pm 0.1$ L | $14.1 \pm 0.3$ L | $15.6 \pm 0.3$ L | $17.8 \pm 0.2$ L |
| | | | | $0.146 \pm 0.003$ CSI |
| | | | | $0.08 \pm 0.003$ RVM |
| Wild | | | | $0.224 \pm 0.005$ CSI |
| | | | | $0.16 \pm 0.006$ RVM |

At the end of the smoltification period, we found a significant effect in the relative ventricular mass (RVM; χ*^2^_(2)_* = 159.4, *p* < 0.001) between all groups. Specifically, wild fish had a significantly lower RVM compared to both SIMNAT and SIMFARM (*p* < 0.001, for both). Furthermore, SIMNAT tended to have a lower RVM than SIMFARM (*p* = 0.09). Similarly, there was a significant effect of treatment on the cardio somatic index (CSI; χ*^2^_(1)_*= 172, *p* < 0.001) at the end of the smoltification period. That is, CSI was higher in wild fish compared to both SIMFARM and SIMNAT (*p* < 0.001, for both), with SIMNAT having lower CSI (*p* < 0.001) than SIMFARM (Table 1).

### Fish reared in SIMNAT develop a heart morphology that resembles that of wild salmon

Heart morphology differed dramatically between the groups at smoltification, but some of these differences appeared already during the fry and parr stages. Firstly, while height:width ratio was not different between groups at the fry stage, it decreased in SIMFARM, compared to wild fish during the parr stage. Furthermore, the height:width ratio was higher in wild compared to both SIMNAT and SIMFARM smolts, indicating that wild fish develop more elongated and narrow ventricles compared to experimental fish, and that rounding of the ventricle develop already during the parr stage in SIMFARM fish. SIMNAT fish tended to have more elongated ventricles than the SIMFARM fish at smoltification (*p* = 0.06), however this difference did not reach statistical significance. Of note, whereas height:width ratio remained stable throughout all sampling points in SIMNAT and wild fish, it decreased in SIMFARM between the fry and the parr stages as well as between fry and smolt stages. Thus, SIMFARM ventricles became gradually more rounded through the freshwater phase (Figure 2A).

**Figure 2.**
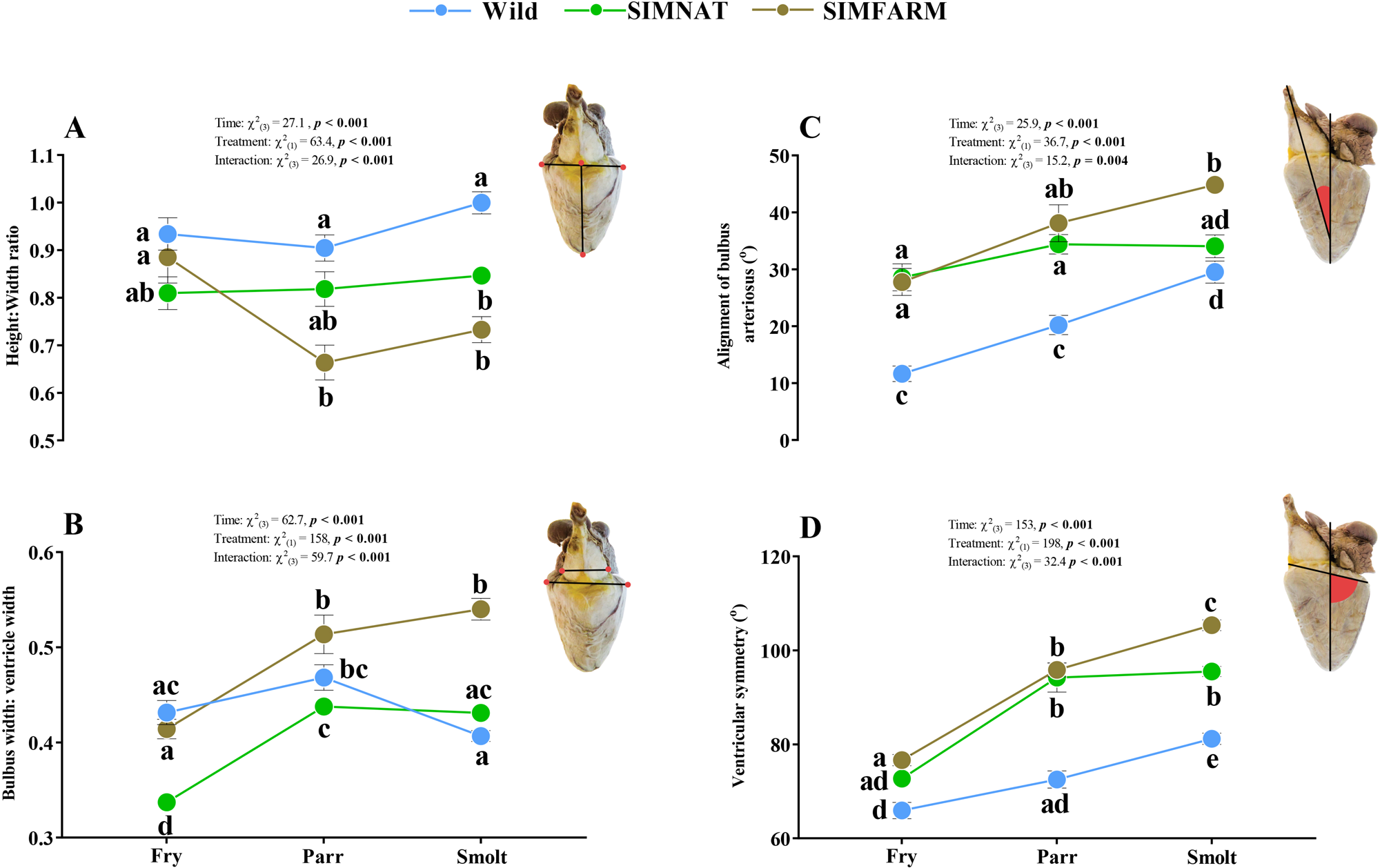
Heart morphology average (± SEM) measurements throughout the freshwater life stages (fry, parr and end of smoltification) for Atlantic salmon reared under simulated farmed (SIMFARM), simulated natural (SIMNAT) conditions or captured from the wild. Linear Mixed Effects model statistics are given in each panel and small letters represent Tukey post-hoc differences between groups at all timepoints.

Bulbus:ventricle width ratio - a measurement of relative bulbus size - increased in both SIMFARM and SIMNAT fish from fry to smolt and was consistently higher in SIMFARM compared to SIMNAT throughout the freshwater phase. Bulbus:ventricle width ratio appeared more stable in wild fish and was higher than in SIMNAT at the fry stage and lower than in SIMFARM fish at smoltification (Figure 2B).

Alignment of the bulbus arteriosus angle was consistently higher in SIMFARM compared to wild fish, but both groups experienced a gradual increase in this angle throughout the period, indicating that the bulbus becomes less aligned with time (Figure 2C). In SIMNAT fish, this angle was larger than in wild fish already at the fry stage but did not increase further throughout the freshwater phase. Thus, at smoltification, this angle was lower in SIMNAT compared to SIMFARM, but there was no difference between SIMNAT and wild.

Finally, the ventricular symmetry angle increased from fry to smolt stage in all groups but ventricles of SIMFARM fish were consistently more asymmetric than those of wild fish (i.e. larger ventricular symmetry angle). Ventricular symmetry was similar in SIMNAT and SIMFARM until the smolt stage where ventricular asymmetry was more pronounced in SIMFARM compared to SIMNAT. Furthermore, ventricular symmetry angle was also higher in SIMNAT compared to wild fish both at the parr and smolt stages (Figure 2D).

All in all, ventricles of SIMNAT fish tended to be more similar to those of wild fish compared to SIMFARM fish.

### Differences between SIMNAT and SIMFARM are already present at the start of the smoltification period

In order to increase our understanding of the temporal resolution of morphological development during the freshwater stage, we included a sampling point for the experimental groups during the start of the smoltification period. While differences in ventricle roundness (*i.e.* height:width ratio) were only evident at the end of the smoltification period, all other measurements were already significantly different (bulbus:ventricle width: *p* < 0.001, alignment of the bulbus arteriosus angle: *p* = 0.04 and ventricular symmetry angle: *p* < 0.001) between SIMNAT and SIMFARM at the start of the smoltification period (Supplementary Figure 3).

### Surface area and volume MRI image analysis

Tissue composition and lumen properties of hearts from wild, SIMNAT and SIMFARM smolt were assessed by MRI. The compactum thickness (normalized to the spongiosum area) was greatest in SIMFARM fish compared to both SIMNAT and wild fish (*p* < 0.001, for both). Meanwhile, relative compactum thickness was not different between wild and SIMNAT fish (*p* = 0.37, Figure 3A).

**Figure 3.**
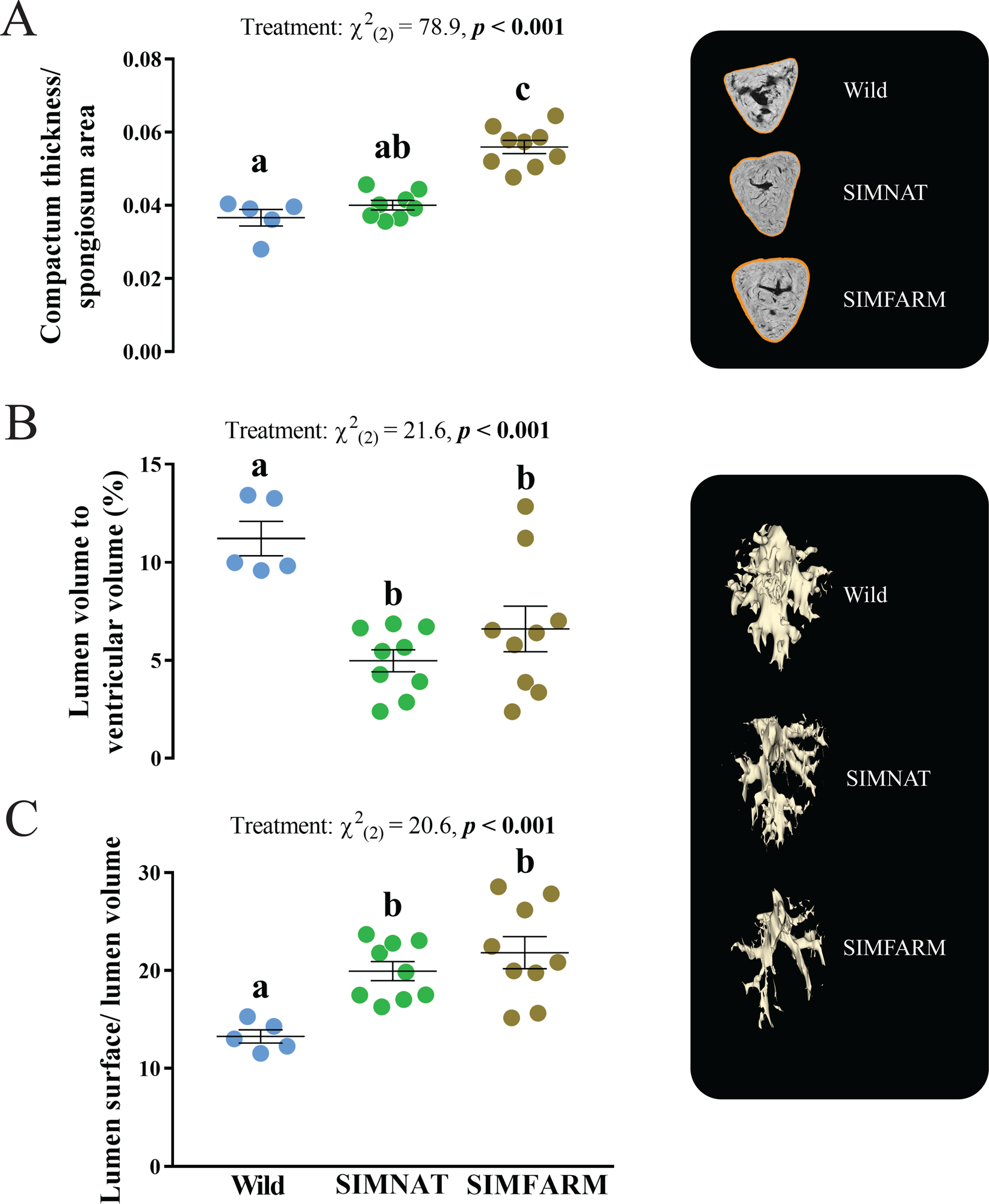
Compactum thickness (normalized to the spongiosum area; **A**), lumen volume as percent of ventricular volume; **B**) and lumen surface area (normalized to lumen volume; **C**) measurements taken at the end of the smoltification period in Atlantic salmon reared under simulated farmed conditions (SIMFARM), simulated natural conditions (SIMNAT) and in the wild (from the Nidelva population in southern Norway). Linear Mixed Effects model statistics are given in each panel and small letters represent Tukey post-hoc differences between groups.

Wild fish had the highest percent of lumen volume (normalized to ventricular volume) and the lowest lumen surface (normalized to lumen volume) compared to both experimental groups (SIMNAT: *p* = 0.001 and *p* = 0.01, respectively and SIMFARM: *p* = 0.01 and *p* = 0.001, respectively). There were no significant differences between SIMNAT and SIMFARM groups in either volume (*p* = 0.4) or lumen surface (*p* = 0.53; Figure 3B and 3C).

### Cardiac function

Systolic and diastolic cardiac function were assessed in 10 SIMNAT and 10 SIMFARM individuals with echocardiography (Supplementary Figure 1). The heart rates for SIMFARM and SIMNAT were 75.9 ± 3.8 and 75.8 ± 6.9 ml/min/kg, respectively.

#### Diastolic function

There was a significant increase (*t*_(17.8)_ = 2.27, *p* = 0.03) in both early, E (Figure 4A) and late diastolic, A (*t*_(10.3)_ = 2.39, *p* = 0.04) blood flow velocities in SIMFARM, compared to SIMNAT (Figure 4B). This parallel increase resulted in unchanged (*t*_(12.9)_ = -0.05, *p* = 0.96) E:A ratios (Figure 4C).

**Figure 4.**
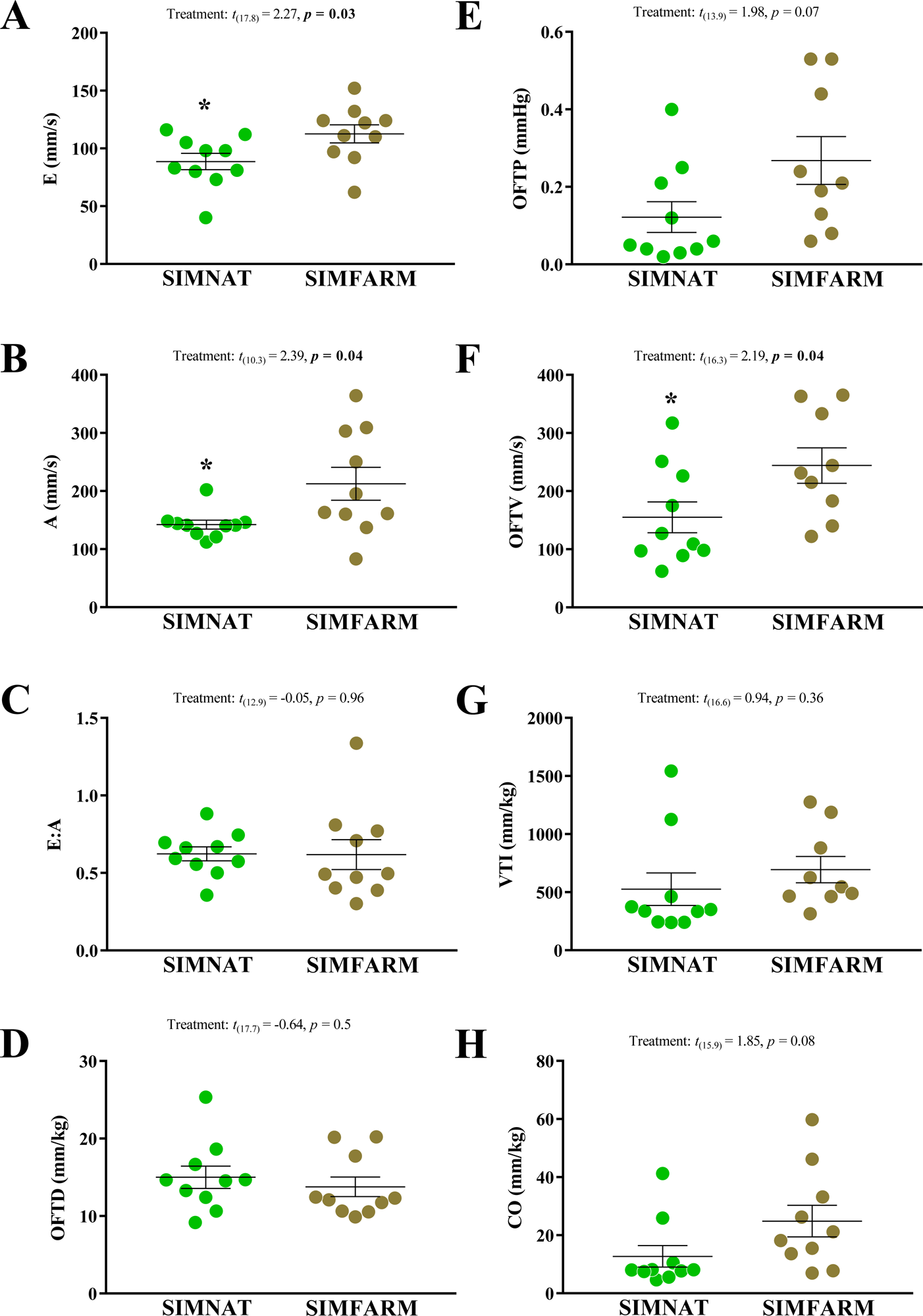
Echocardiography measurements of the early diastolic atrio-ventricular valve velocity (E; **A**), late diastolic atrio-ventricular valve wave velocity (A; **B**), the ratio between the E and A (EA; **C**), the outflow tract diameter (**D**), outflow tract pressure (**E**), outflow tract velocity (**F**), blood velocity time integral (**G**) and cardiac output (**H**), taken at the end of the smoltification period in Atlantic salmon reared under simulated farmed (SIMFARM) or simulated natural (SIMNAT) conditions. Statistics are given in each panel and * represent significant differences between groups.

#### Systolic function

There were no significant differences (*t*_(17.7)_ = -0.64, *p* = 0.5) in the outflow tract diameter (Figure 4D), but a tendency towards increased (*t*_(13.9)_ = 1.98, *p* = 0.07) pressure gradients across the VB-valve in SIMFARM, compared to SIMNAT (Figure 4E). Maximal outflow tract blood velocities (*t*_(16.3)_ = 2.19, *p* = 0.04) were also higher in SIMFARM individuals (Fig. 4F) whereas blood velocity time integral was not significantly different between groups (*t*_(16.6)_ = 0.94, *p* = 0.36). Consequently, cardiac output (*t*_(15.9)_ = 1.85, *p* = 0.08) was similar between the groups (Figure 4H).

Correlation analyses of morphological and functional parameters were performed to understand the relationship between morphology and function (Figure 5). We observed a higher CSI accompanied by a tendency for higher relative ventricular mass in SIMFARM. MRI recordings showed that this appears to be caused by hypertrophied ventricular compact tissue. Indeed, the RVM correlated positively with the outflow tract pressure (ρ = 0.64, *p* = 0.02) and the outflow tract velocity (ρ = 0.65, *p* = 0.02), and tended to positively correlate also with the blood velocity time integral (ρ = 0.51, *p* = 0.09) and CO (ρ = 0.54, *p* = 0.08). Meanwhile, CSI tended to positively correlate with the outflow tract pressure (ρ = 0.56, *p* = 0.07), the outflow tract velocity (ρ = 0.57, *p* = 0.07), the blood velocity time integral (ρ = 0.5, *p* = 0.09), the cardiac output (ρ = 0.54, *p* = 0.08), A (ρ = 0.52, *p* = 0.09), as well as with the bulbus:ventricle width ratio (ρ = 0.57, *p* = 0.07). In addition, the RVM was positively correlated with A (ρ = 0.65, *p* = 0.02) and tended to correlate positively also with E (ρ = 0.53, *p* = 0.08). Furthermore, the bulbus:ventricle width ratio tended to correlate positively with the ventricular symmetry angle (ρ = 0.51, *p* = 0.09). Finally, we found a tendency for a negative correlation (ρ = -0.54, *p* = 0.08) between the height:width ratio and the E:A ratio.

**Figure 5.**
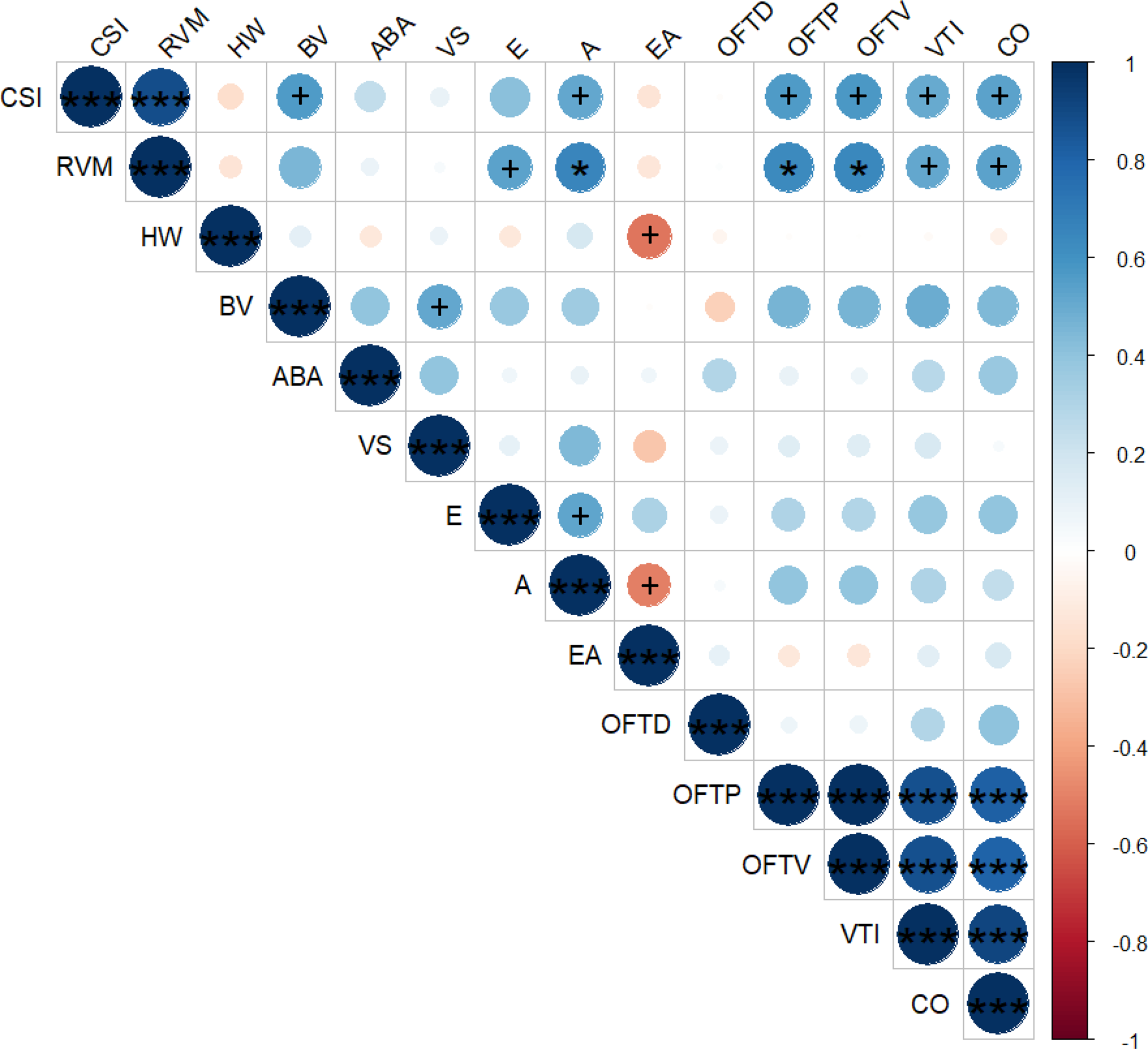
Spearman’s correlation matrix with Benjamini-Hochberg *p* value correction for multiple comparisons between the cardiosomatic index (CSI), the relative ventricle mass (RVM), the ventricular height:width ratio (HW), the bulbus:ventricular ratio (BV), the alignment of the bulbus arteriosus (ABA), the early diastolic atrio-ventricular valve velocity (E), the late diastolic atrio-ventricular valve wave velocity (A), the ratio between the E and A (EA), the ventricular symmetry (VS), the outflow tract (OFT), the outflow tract pressure (OFTP), the outflow tract velocity (OFTV), the blood velocity time integral (VTI) and the cardiac output (CO). Positive and negative correlations are illustrated with a blue and red gradient (respectively) and the magnitude of the correlation is indicated by the size of the circles within each square. Adjusted significant correlations (p < 0.05) are indicated by * and tendency for significance (0.05 > p < 0.1) are indicated by a + symbol.

## Discussion

In this study, we demonstrate the pivotal role of environmental signals (*i.e.* photoperiod and temperature) in shaping heart morphology during early rearing stages of Atlantic salmon. Specifically, our findings highlight that rearing conditions mimicking a natural environment (SIMNAT), characterized by colder water temperatures and a more natural light regime, facilitate the development of an elongated, pyramid-like shaped heart with a symmetrical structure, closely resembling wild salmon heart. In contrast, simulated farmed (SIMFARM) rearing conditions are associated with the development of a heart with rounded, asymmetrical morphology that deviates significantly from the wild salmon heart. This distinct cardiac morphology has previously been linked to impaired cardiac and physical performance (Claireaux et al., 2005), which is in agreement with our results. Specifically, we found that the morphology observed in SIMFARM is associated with increased blood flow velocities in the bulbus, which implies a higher cardiac workload. Our results underscore the significance of environmental factors during early life stages as fundamental drivers of this geometrically advantageous heart shape. Further, we observed that specific morphological differences between our treatment groups emerged early on and persisted throughout the experiment. Conversely, other variations appeared later, suggesting that different environmental factors may be time-specific in their impact on cardiac morphological traits.

At smoltification, heart morphology differed dramatically between SIMNAT and SIMFARM salmon, with SIMNAT hearts being more similar to wild salmon smolt. For example, the ventricles of SIMNAT fish were more elongated (higher ventricular height:width ratio) than those of SIMFARM fish but rounded compared to those of wild smolt. These results agree with previous findings showing that sea farmed Atlantic salmon and rainbow trout have rounded ventricles compared to wild salmon (Engdal et al., 2024; Poppe et al., 2003). Moreover, in agreement with our data, rounded ventricles have previously been associated with elevated rearing temperatures during early life stages in salmonids (Brijs et al., 2020; Frisk et al., 2020). Notably, unlike their wild counterparts, farmed salmon appear to develop increasingly rounded ventricles as they progress through their life cycle. Specifically, we found that mean height:width ratios of SIMNAT fish were consistently lower, than those observed in wild salmon.

Nonetheless, these ratios remained stable throughout the freshwater stage, while those of SIMFARM fish exhibited a different trend. That is, already at the fry stage SIMFARM fish had lower height:width ratios (~0.9) than SIMFARM and wild which decreased even further (~0.6-0.7) at the parr and smolt stage. These differences and changes in ventricle roundness indicate not only that the environmental conditions have a dramatic effect on ventricular shape, but that environmental conditions in the earliest rearing stages (*i.e.* before fry stage) are probably important. In the aquaculture industry, it is common to use rearing temperatures far exceeding those experienced by developing embryos in nature (Handeland et al., 2003) and future research should examine the impact of rearing temperature during embryogenesis on ventricular shape and function. Importantly, cardiac roundness has previously been associated with impaired cardiac and physical performance in rainbow trout (Claireaux et al., 2005) as well as increased mortality risk in Atlantic salmon (Frisk et al., 2020).

Using echocardiography, we found that the morphology observed in SIMFARM is associated with increased blood flow velocities in the bulbus. In addition, both E and A wave velocities were elevated during ventricular filling in this group, and there was a tendency for a negative correlation between the E:A ratio and ventricular height:width, indicating that rounder ventricles display decreased diastolic function. Typically, faster blood flow during both diastole and systole are associated with concentric hypertrophy in situations where the ventricular outflow tract is partially obstructed (Agatston et al., 1989; Kenny, Plappert and St John Sutton, 1991). Indeed, significant correlations were observed between relative heart and ventricle weight and both diastolic and systolic blood flow velocities, supporting that larger (hypertrophied) hearts produce faster blood flow. In addition, we found that SIMFARM fish exhibited significantly enlarged bulbi in comparison to both SIMNAT and wild fish at the end of the smoltification period and it remained higher in SIMFARM from parr to smolt. The enlargement of the bulbus arteriosus could imply alterations in the elastic properties of this cardiac chamber. That is, the bulbus arteriosus is part of the outflow tract of the teleost heart and extends from the ventricle to the ventral aorta delivering blood to the gills. To protect the gill capillaries from high systolic pressure, the bulbus is a compliant chamber that serves to steady pulsatile blood flow across the capillary networks of the gill (Evans, Claiborne and Currie, 2013). In other words, if the enlarged bulbi are also less compliant, this may create and obstruction leading to increased blood flow velocities. We also observed that fish reared under simulated farmed conditions develop more misaligned bulbi compared to wild fish (Engdal et al., 2024; Poppe et al., 2003). Interestingly, Brjis et al. (Brijs et al., 2020) observed that the ability of rainbow trout to tolerate confinement stress was correlated with a lower bulbus arteriosus angle alignment, which suggests that this angle may play a role in cardiac stress resilience. A misaligned and less compliant bulbus is thus expected to increase the load on the heart. When the heart faces increased workload, due to either physiological or pathological factors, it responds by hypertrophic growth of the compact myocardium (Graham and Farrell, 1992; Johansen et al., 2011; Johansen et al., 2017; Keen et al., 2016). An increase in the thickness of the compact myocardium may therefore represent an adaptive response to environmental changes (Graham and Farrell, 1992; Keen et al., 2016) or a pathological reaction to cardiac overload (Johansen et al., 2011; Johansen et al., 2017). No large differences were observed in fish performance (growth, mortality, etc.) at these early life stages, but it is tempting to speculate that the morphological and functional differences observed in this study may lead to cardiac disorders later in life. However, further studies are needed to clarify this.

To the best of our knowledge, lumen volume and surface area data have not been previously reported for any teleost fish species. In our current study, we employed high-resolution MRI scans to quantitatively assess these parameters in both experimental and wild salmon. While lumen volume and surface area remained comparable between the two experimental groups (SIMFARM and SIMNAT), the substantial disparities observed between wild and experimental fish warrant further consideration. Notably, the ventricular lumen volume in experimental smolt (comprising both SIMFARM and SIMNAT groups) was less than half that of their wild counterparts. This drastic reduction in lumen volume appears directly associated with a substantial, albeit not proportionate, increase in lumen surface area in the same fish. Given that measurements of these lumen characteristics have not been previously documented, we are limited to speculation regarding their underlying causes and significance. However, we speculate that an augmentation in lumen surface area may potentially arise from increased branching of the spongy myocardium. This tissue relies solely on venous blood returning to the heart for oxygenation and the increased surface area may signify an elevated demand for oxygen or nutrients. This, however, likely contributes to a reduction in lumen volume and possibly enhances blood turbulence within the heart. It is difficult to imagine how this remodeling would be inherently adaptive, except through providing increased surface area for enhanced resource uptake. If this holds true, it implies that experimental fish experienced specific stimuli in their rearing conditions, necessitating such compensatory remodeling. While this finding is undoubtedly fascinating and merits careful consideration, it is beyond the scope of this article to speculate on the collective influences of various rearing conditions on the observed differences in internal cardiac characteristics.

Despite our efforts to mimic natural conditions for the SIMNAT fish, rearing conditions for wild and experimental fish differed considerably. Experimental fish were confined indoors in controlled tank environments, with reduced possibility for physical activity and unlimited access to food, in stark contrast to the conditions of their wild counterparts. This was also reflected in morphological differences between wild and SIMNAT fish. These differences may have been associated with other factors affecting cardiac remodeling. For example, the elongated pyramid-shaped ventricle appears to be adapted for physical activity, and thus increased training may play a crucial role in cardiac shape. However, further investigations are needed to explore this aspect.

## Conclusions

The SIMFARM phenotype in this study is characterized by a rounded ventricle, a thicker compactum and an increased misalignment of the bulbus arteriosus, compared to both the SIMNAT and the wild fish. These morphological differences seem to be associated with increased cardiac workload and diastolic dysfunction. However, more comprehensive studies of the potential implications of altered cardiac morphology on cardiac function are needed. Importantly, our findings have implications reaching beyond the industrial farming of salmonids. Our results are in agreement with other studies which report that elevated temperatures impact heart morphology in a manner that is associated with impaired cardiac and physical performance (Brijs et al., 2020; Frisk et al., 2020). These findings highlight the potential risk posed by climate change on the fitness of wild salmon populations. Thus, more research should be conducted to understand how and why the environment promotes such morphological remodeling of the heart.

## Acknowledgements

We are grateful to Ivar Helge Matre and the technical staff at the Matre Research station for excellent animal husbandry. As well as Lili Zhang (Institute for Experimental Medical Research, Oslo University Hospital, University of Oslo) for providing expert advice for MRI sample preparation and scanning protocols. protocols, and Silje Marie Ulset and Elise Thovsland Sannes for their help collecting wild smolts.

## Author Contributions

Conceptualization: Ø.Ø., O.F., I.B.J., Methodology: M.A.V., V.A.E., S.K., M.F., I.B.J., Validation: M.F., I.B.J., Formal Analysis: M.A.V., V.A.E., S.K., M.F., I.B.J., Investigation: M.A.V., V.A.E., E.H., S.K., M.M., M.F, I.B.J., Resources: E.H., O.F., M.F., I.B.J., Writing - Original draft: I.B.J., M.A.V., V.A.E., Writing - Review and Editing: M.A.V., V.A.E., E.H., Ø. Ø., S.K., M.M., M.F., I.B.J., Visualization: M.A.V., V.A.E., S.K., Project Administration: I.B.J., M.A.V., O.F., Supervision: I.B.J., M.F., Funding Acquisition: I.B.J., Ø.Ø., M.F.

## Data accessibility

All relevant data are within the paper and its Supporting Information files.

